# Sex-biased migration and demographic history of the big European firefly *Lampyris noctiluca*

**DOI:** 10.1101/2024.01.24.577017

**Authors:** Ana Catalán, Daniel Gygax, Ulrika Candolin, Sergio Tusso, Pablo Duchen, Sebastian Höhna

## Abstract

Differential dispersion between the sexes can impact population structure and connectivity between populations, which in turn, can have an effect on the evolvability capability of a species. Here we explored the demographic history of the big European firefly, *Lampyris noctiluca*, which exhibits female neoteny. *L. noctiluca* distribution extends throughout Europe, but nothing is known on how its colonization processes. To investigate this, we produced the first *Lampyris* genome (653Mb), including an IsoSeq annotation and the identification of the X chromosome. We collected 115 individuals from six populations of *L. noctiluca* (Finland to Italy) and generated whole genome re-sequencing for each individual. We inferred several population expansions and bottlenecks throughout the Pleistocene that correlate with glaciation events. Surprisingly, we uncovered strong population structure and low gene-flow. We reject a stepwise, south to north, colonization history scenario and instead uncovered a complex demographic history with a putative eastern European origin. Analyzing the evolutionary history of the mitochondrial genome as well as X-linked and autosomal loci, we found evidence of a maternal colonialization of Germany, putatively from a western European population, followed by male-only migration from south of the Alps (Italy). Overall, investigating the demographic history and colonization patterns of a species should form part of an integrative approach of biodiversity research. Our results provide evidence of sex-biased migration which is important to consider for demographic, biogeographic and species delimitation studies.

## Introduction

Fireflies are a diverse group of bioluminescent beetles with a wide distribution around the globe (Lewis et al. 2020). In Central Europe, around 38 firefly species have been reported (www.gbif.org), for which genomic resources and information about their demographic history is completely lacking. From all 38 species, *Lampyris noctiluca*, (the big European firefly or common glow-worm) has a specially wide distribution across Europe (**Figure 2A**), inhabiting different types of ecological niches (De Cock 2009; Novak 2018). Its wide distribution makes this species particularly interesting to understand past migration patterns in Europe and present insect connectivity across highly fragmented landscapes. *L. noctiluca* adult females are neotenic, retaining a larval-like morphology, a characteristic that is present in several firefly species (Bocakova et al. 2007; South et al. 2011). Short after eclosion adult females start signaling by producing bioluminescent signals to attract males and die after laying eggs (Novak 2018; Van den Broeck et al. 2021). Thus, short-distance migration can take place in both sexes during the larval stage (1-2 years), but in adulthood only winged males disperse (Lehtonen et al. 2021). *L. noctiluca*’s current distribution is marked by the last ice age events, making northern latitudes and in high-altitude locations only available for colonization after the retreat of the glaciers. Range expansion in neotenic insects such as *L. noctiluca* is limited by the movement of the larvae (Lehtonen et al. 2021). Migration between populations is unlikely for the larvae, impossible for adult females, but potentially feasible for adult males. Male-biased migration can produce particular nucleotide diversity patterns, such as nuclear-mitochondria tree discordance and can reduce nucleotide diversity levels when compared to species where gene flow is enabled by both sexes. Currently, we have almost no information about the impact of sex-specific neoteny on population genetic patterns (Eberle et al. 2019) or about the putative consequences that sex-specific neoteny can have in adaptive processes.

The current genetic diversity observed in a species is the product of its past demographic and selection events and can determine its capacity to evolve. Specifically, the last ice age set boundaries on migration patterns which, together with ecological factors, have defined today’s species distribution. The study and discernment of present-day genetic variation are crucial to understand a species susceptibility to climate change. In fruit flies and humans, the various forces shaping genetic diversity have been studied in detail, resulting in well-founded hypothesis for population changes through time, gene flow and adaptation events (Stephan and Li 2007a; Gutenkunst et al. 2009; Pool et al. 2012). This scenario is different for species with no obvious health or economic importance to humans, as is the case for may insects, but see (Catalan et al. 2022).

Despite the importance of insects for their diverse roles in ecosystem function, their high diversity in species number and natural histories, has challenged in some way, the generation of detailed knowledge of their ecological and evolutionary status. Nevertheless, this knowledge is highly relevant to the maintenance and productivity of diverse ecosystems, particularly in the face of the current biodiversity crisis. Astonishingly, most insect taxa lack the necessary genomic resources needed to generate comprehensive demographic and phylogeographic hypothesis, which are crucial for conservation purposes.

In this work we sampled a total of 115 individuals from six populations of *L. noctiluca*, ranging from Finland to Italy and generated population level whole genome re-sequencing data. The goal of this study was to generate the first demographic hypothesis, including population size changes through time and migration events for this species with a particular focus on the impact of female neoteny. Additionally, we assembled the first genome for *L. noctiluca*, contributing in this way to the genomic resources for fireflies and insect research. We investigated several population genetic statistics and generated hypothesis for *L. noctiluca*’s colonization patterns by taking advantage of methods such as StairwayPlot2 (Liu and Fu 2015), PoMo (<u>Po</u>lymorphism-aware phylogenetic <u>Mo</u>dels) (Borges et al. 2022) and ABC demographic modeling. We uncovered a complex demographic history, putatively with multiple migration routes from different glacial refugia and identified genetic footprints left by sex-biased migration. Our work contributes to generate a more complete vision of migration patterns in Europe and marks the first step into generating an integrative understanding of the forces shaping genetic diversity in fireflies.

## Results

### Genome assembly

We generated long Nanopore and short Illumina reads to assemble a genome for *L. noctiluca* (*LaNoc*) (**Figure 1**). From the four explored assembly strategies (MaSuRCa, Flye, Canu and Shasta) (Zimin et al. 2013; Kolmogorov et al. 2019; Shafin et al. 2020), the hybrid approach with MaSuRCA gave the best results (**Table S1**). LaNoc’s genome lies at the lowest quantile of known genome sizes for fireflies (Lampyridae) (Liu et al. 2017; Lower et al. 2017) (**Figure 1A**) with 652,498,014 bp (**Table 1**). We identified the X chromosome using a female-to-male difference in genome coverage strategy, successfully detecting contigs with half coverage values in males, which is the heterogametic sex in fireflies (X0 or XY) (Wasserman and Ehrman 1986; Dias et al. 2007). The X chromosome is 19 Mbp in length, comprising ∼2.9% of the genome (**Figure 1C**). We identified a complete mitochondrial genome covering 18,937 bp. Based on PacBio IsoSeq RNA sequencing, we annotated ∼31K transcripts and delimited genomic regions into transcripts, exons, introns and intergenic. Repetitive elements comprised 41.76% of the genome, most of which are unclassified repeats (18.16%), followed by retroelements (12.41%) and DNA transposons (10%) (**Figure 1D**).

**Table 1.** General genome assembly statistics.

| Statistic | Value |
| --- | --- |
| Genome size (bp) | 652,498,014 |
| No. of contigs | 1,401 |
| No. IsoSeq transcripts | 31,648 |
| Largest contigs | 4,636,015 |
| Chr X length (bp) | 19,305,488 |
| Mitochondria (bp) | 18,937 |
| N50 (bp) | 655,439 |
| L50 | 292 |
| CG % | 35.17 |
| Repeat content | 41.76% |
| Genome heterozygosity | 0.91% |
| BUSCO Insecta % | 94.60 |

**Figure 1.**
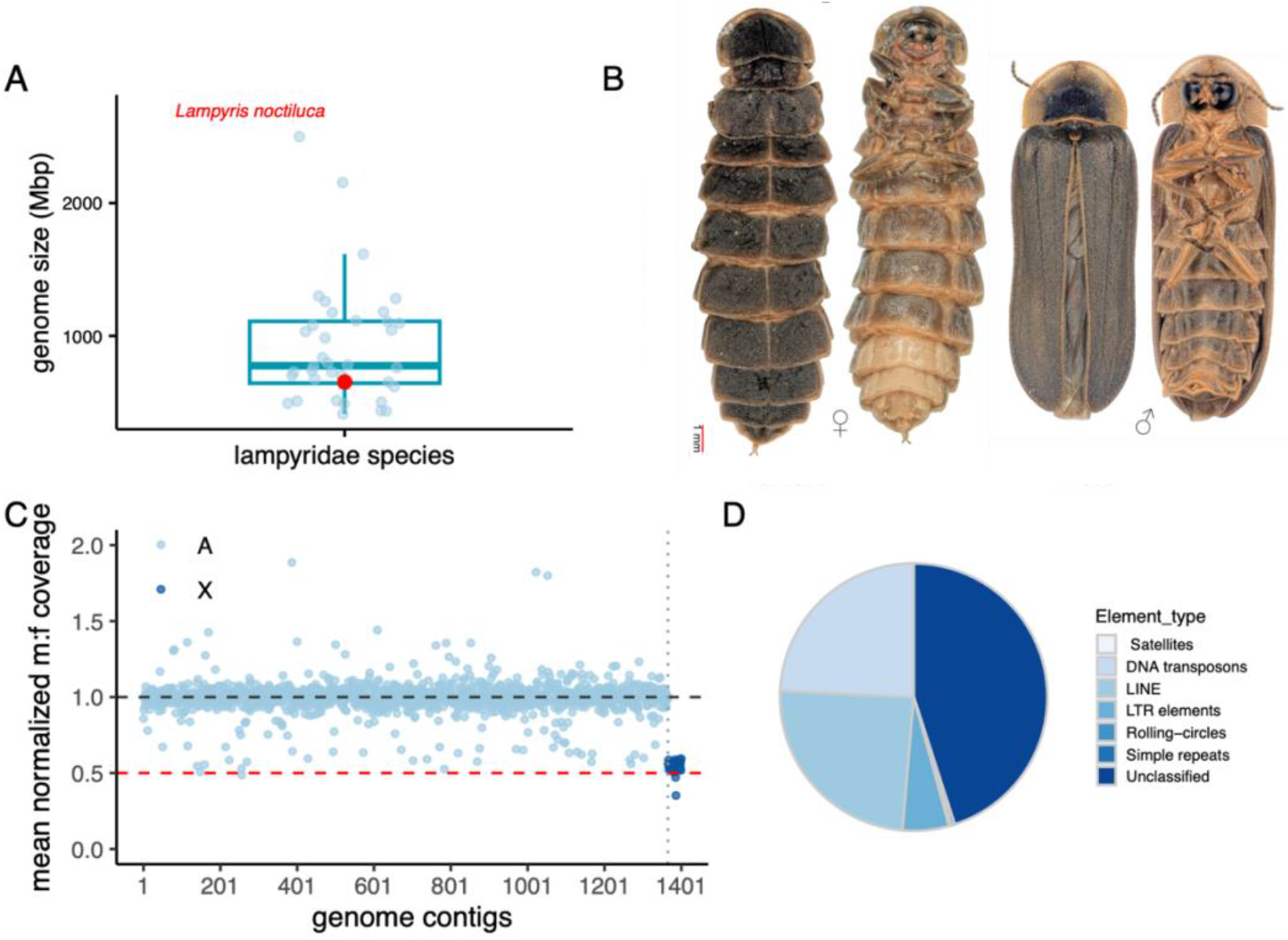
Genome sequencing of *Lampyris noctiluca*. (**A**) Boxplot showing genome size distribution of available fireflies including genome size of *Lampyris noctiluca* highlighted in red. (**B**) Dorsal and ventral view of female (left) and male (right) images of *L. noctiluca*. (**C**) Male-to-female ratio of normalized genome coverage across all assembled contigs. Dark blue dots correspond to X chromosome linked contigs, i.e., contigs with a significantly lower m:f ratio than 1, tested by a Wilcoxon-test (**Table Sxx**). (**D**) Piechart showing percentages of repetitive element types in the genome. Repeats constitute 41.76% of the genome.

**Figure 2.**
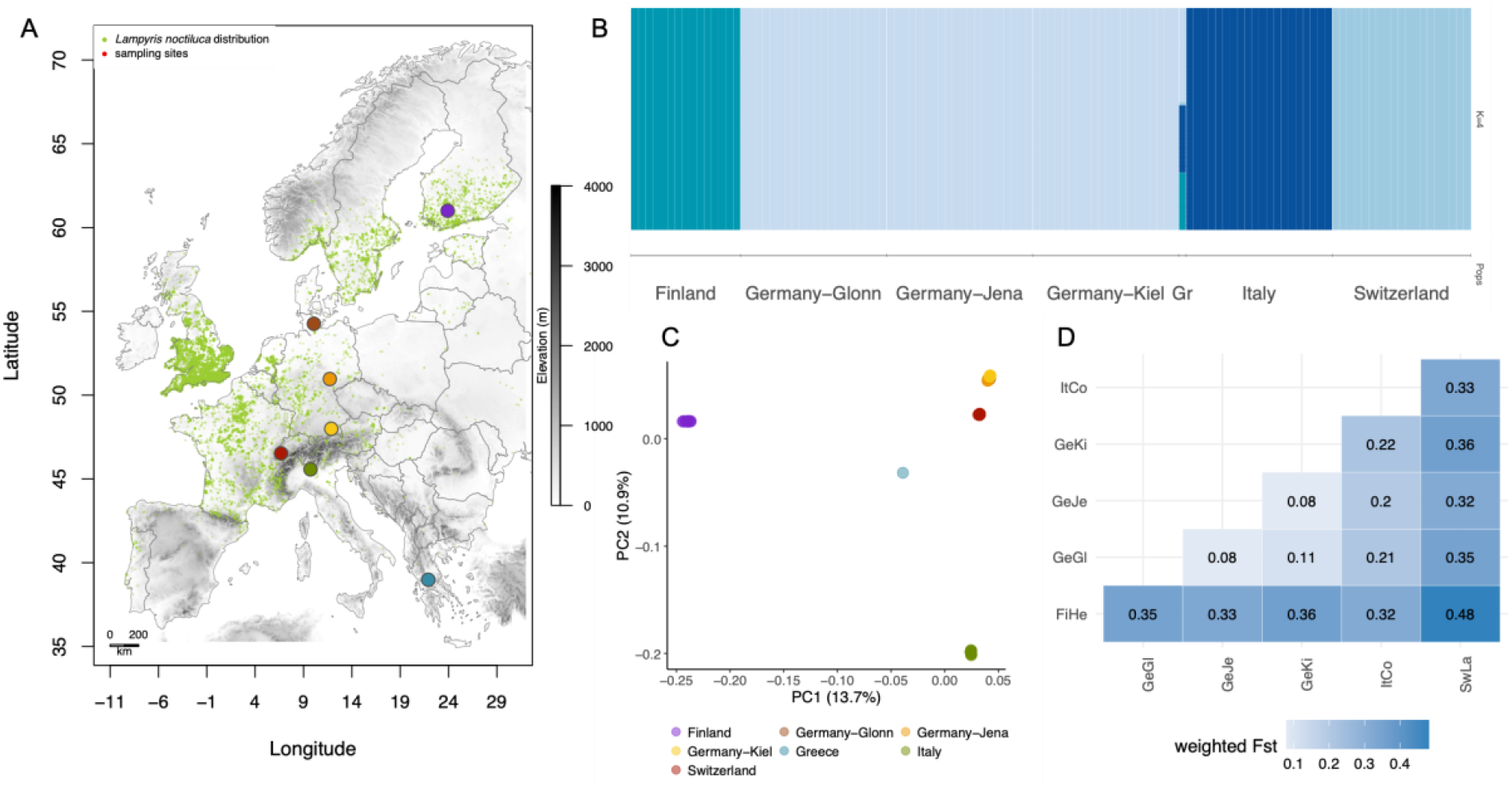
Population structure in *L. noctiluca*. (**A**) Map of Europe marking the locations of sample collection. (**B**) Genetic clusters assessed with STRUCTURE (k=4). (**C**) Principal component analysis (PCA). (**D**) Pairwise population *Fst* statistic for population differentiation. FiHe: Finland, GeKi: Germany-Kiel, GeJe: Germany-Jena, GeGl: Germany-Glonn, ItCo:Italy, SwLa: Switzerland.

### Population structure

We sampled a total of six *L. noctiluca* populations from Finland, Germany-Kiel, Germany-Jena, Germany-Glonn, Switzerland, Italy and one outgroup specimen collected in Greece, sequencing a total of 115 individuals (**Figure 2A**). We identified 61,441,736 autosomal and 841,085 X chromosome SNPs across all individuals. Genetic clustering approaches, such as PCA and STRUCTURE, showed well defined populations that clustered by country (**Figure 2B, C, Figure S1**). The three German populations are very closely related to each other, mostly forming one single genetic cluster, although the ADMIXTURE analysis at k=4 and k=5 revealed some level of population substructure (**Figure S2**). Population differentiation (*Fst*) (**Figure 2D**) and population divergence (*Dxy*) (**Figure S3**) were calculated for every population pair, with both statistics showing of strong population differentiation (**Figure 2D**). Overall, these results show strong population structure, which could be the result of limited migration, as mobilization only occurs at the larval stage and by flying adult males.

### Past population size changes

Changes in effective population size over time is an intrinsic characteristic of populations and provides information about past events. We used a Bayesian variant (implemented in the software RevBayes) of the StariwayPlot 2 (Liu and Fu 2020) approach to uncover single population demographic histories. All populations showed pronounced changes in population size over the last 100k years (**Figure 3A**). The impact of these changes through time were also detected with Tajima’s *D* (Tajima 1989a) (**Figure S4**). Over the last 100k years, the effective population sizes varied from a maximum of 6.3 million individuals (Italy) to a minimum of 7k individuals (Switzerland and Finland). At least two deep bottleneck events were recovered for each population, however, the timing of these bottlenecks varied between populations (**Figure 3A**). For the German populations, however, the timing of the bottlenecks overlap, a pattern that could be driven by common environmental stressors leading to a bottleneck or through a common ancestral population (Stroeven et al. 2016; Seguinot et al. 2018). Towards the present, most populations, with the exception of the Kiel population, show a decline in effective population size. The observed decline in the last 10,000 years can be explained by the colonization of new habitats after the retreat of the last ice age resulting in a population range expansion. Alternatively, it could also be explained by strong selective pressures that resulted in a population decline.

**Figure 3.**
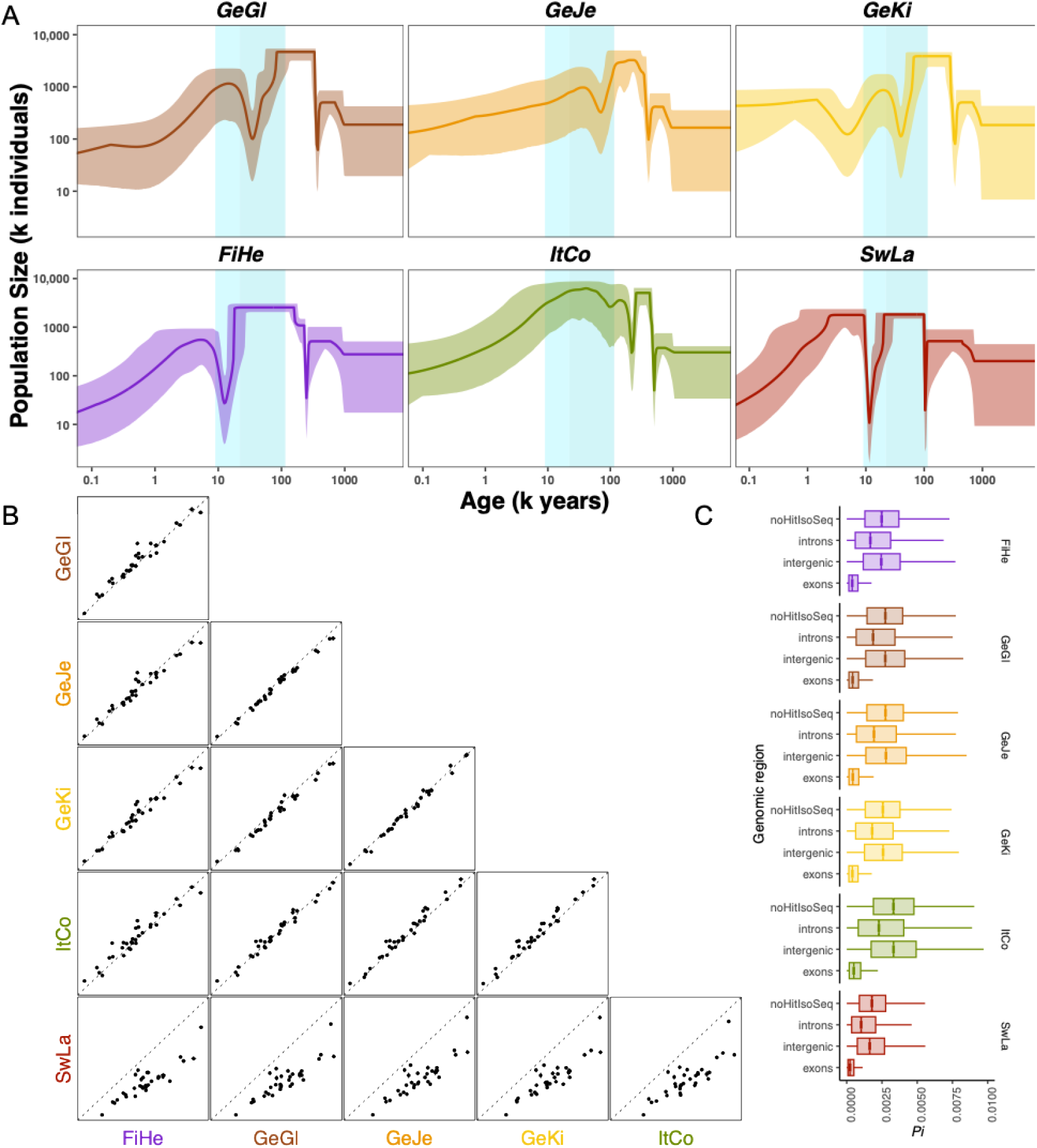
Population size changes past in time. (**A**) StairwayPlot 2 estimation of changes in past population size. Lighter color represents 95% confidence intervals. Light blue vertical rectangle marks the start and the end of the ice age (9000 - 115000 years ago). (**B**) Pairwise nucleotide diversity estimate (*π*) (**C**) Estimation of coalescent ages (most recent common ancestor; MRCA) of all sampled individuals of a population for 35 autosomal. FiHe: Finland, GeKi: Germany-Kiel, GeJe: Germany-Jena, GeGl: Germany-Glonn, ItCo:Italy, SwLa: Switzerland.

### Inference of the time of most recent common ancestor

In the scenario of a species undergoing a range expansion, leading to the colonization of new habitats, we expect to find a shared common ancestor between all derived populations (Excoffier et al. 2009). Conversely, if the populations have been well established long enough ago, then we expect independent common ancestor for each population, i.e., younger than the population splitting time. To explore such a scenario, we inferred the most recent common ancestor (MRCA; coalescent time) for all sampled individuals of a population, repeating the inference for each contig and each population. The inferred MRCAs for each population (with the exception of the Swiss population) was ∼800 kya for the autosomes and ∼350 kya for the X chromosome (**Figure S5**). Such an old time for the MRCA is caused by large population size (see above). The time of the MRCA for each homologous contig was highly correlated across populations, but across contigs the time varied significantly (**Figure 3B**), which provides evidence that a common ancestor was shared among populations. The younger time of the MRCA of the Swiss population, which is also reflected in its lower *π* values (**Figure 3C**), might be explained by a more recent founder event, a stronger bottleneck or smaller effective population size. The relatively old and shared common ancestor is also reflected in the high number of shared ancestral polymorphisms between the populations (**Figure S6**). Overall, the shared coalescent time provides evidence for a common ancestry among the collected populations of *L. noctiluca* (**Figure 3b**).

### Population colonization hypothesis

We first explored a stepwise colonization process from south to north (Greece → Finland) as has been observed in humans, the fruit fly and other firefly species (Stephan and Li 2007b; Excoffier et al. 2013; Catalan et al. 2022). We did not observe a declining pattern of nucleotide diversity (*π*) with increasing latitude (**Figure 3C**) and an isolation by distance scenario was rejected by a Mantel test (*p*-value= 0.137, **Figure S7**). The population’s tree topology, inferred by a polymorphism aware phylogenetic approach (PoMo) (De Maio et al. 2015; Borges et al. 2022), does not depict a south to north colonization pattern (**Figure 4A**). Instead, the PoMo tree showed an unexpected topology, where Greece (diverged >80,000 ya) and Finland (diverged ∼38,000 ya) show the deepest divergence times. Falling into the ice age period, the Swiss and the Italian populations, show divergence times of ∼24,000 and ∼21,000 ya respectively. The three German populations diverged 8,000 to 10,000 years ago, hinting a post ice age colonization. The inferred divergence times and tree topology open up the question of the colonization process of these populations. In the case of the Italian population, it is plausible that it survived in a glacial refugia, as north Italy was not covered by ice during this period (Patton et al. 2017). On the other hand, Finland and Switzerland were covered by glaciers at the estimated divergence times (Becker et al. 2016; Stroeven et al. 2016). The presence of an unsampled or “ghost” population could explain the deep divergence times retrieved for Finland and Switzerland.

**Figure 4.**
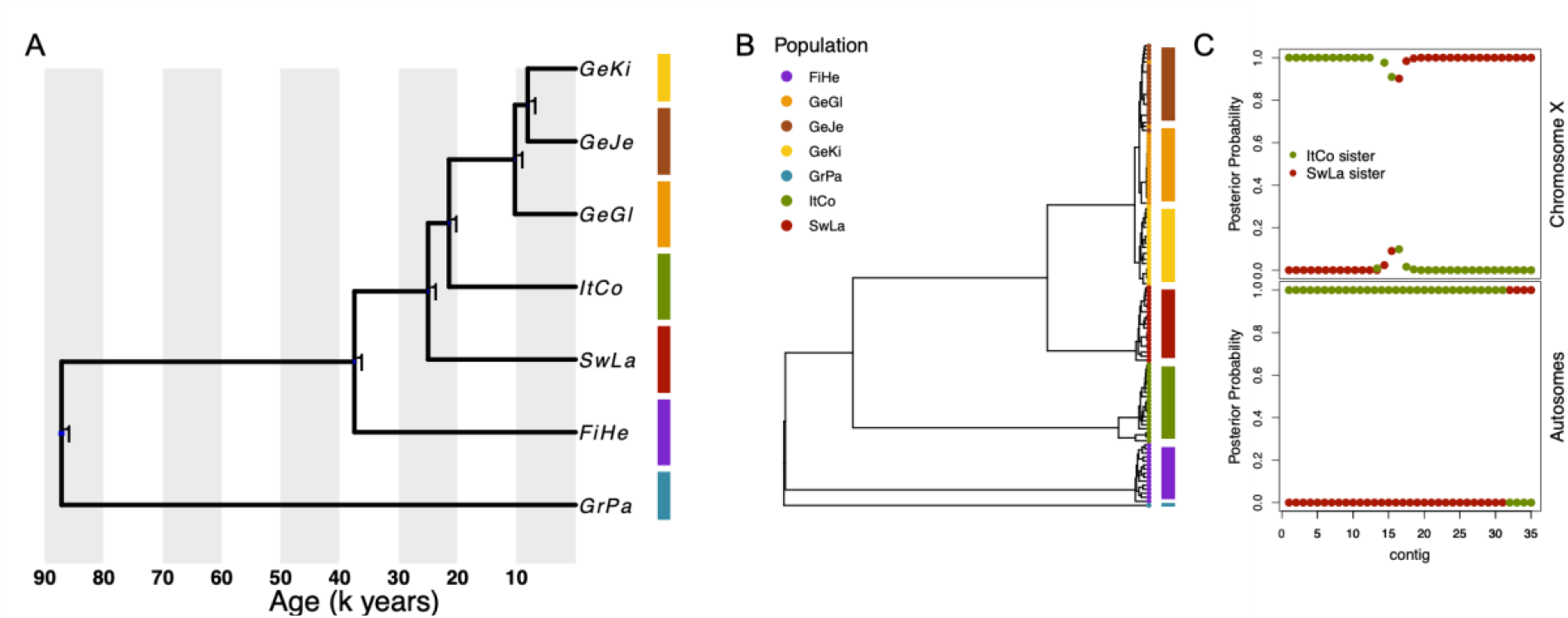
PoMos phylogenetic analysis and nuclear-mitochonria-X tree discordance. (**A**) Genetic relationship across populations estimated by a PoMos analysis. (**B**) Maximum likelihood phylogenetic tree of the mitophondria. (**C**) PoMos posterior probability of either Italy or Switzerland being sister to Germany. FiHe: Finland, GeKi: Germany-Kiel, GeJe: Germany-Jena, GeGl: Germany-Glonn, ItCo:Italy, SwLa: Switzerland, GrPa: Greece-Paleokastro.

**Figure 5.**
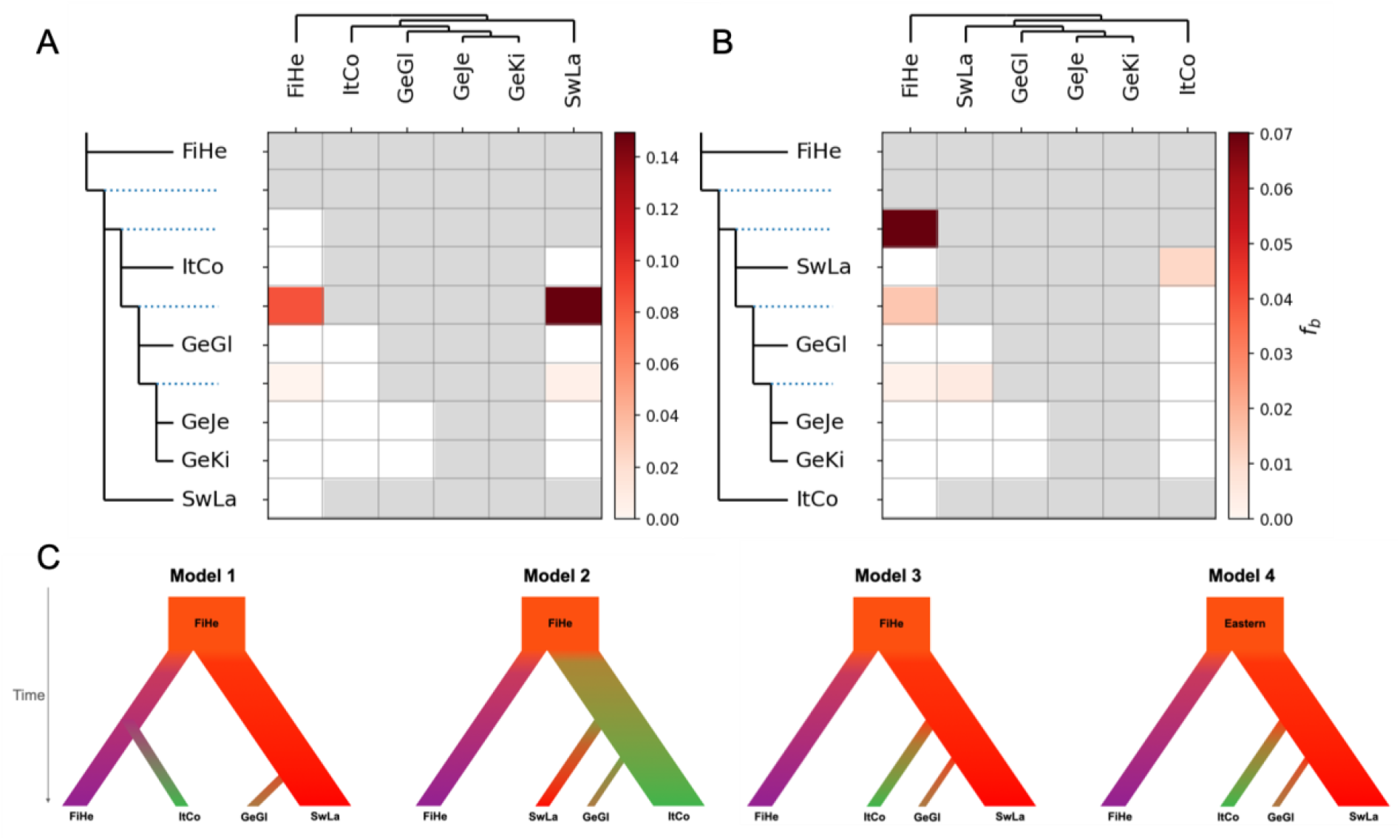
*f*-branch statistics and ABC modelling. (**A**) *f-branch* statistic investigating autosomal migration. (**B**) *f-branch* statistic investigating mitochondrial migration. Gradient bar shows the fraction of shared alleles between tips and branches. (**C**) Tested demographic scenarios presented. *Model 1*: Finland as the most ancestral population from with Switzerland and Italy derive independently. Switzerland is sister to Germany. *Model 2*: Finland as the most ancestral population from which Italy derives. Switzerland and Germany derive from the Italian population. Italy is sister to Germany. *Model 3*: Finland as the most ancestral population from which Switzerland derives. Italy and Germany derive from the Swiss population. Switzerland is sister to Germany. *Model 4*: A ghost population (a population from the East) represents the most ancestral population, from which Finland and Switzerland derive independently. Italy and Germany derive from the Swiss population and Switzerland is sister to Germany.

### Assessment of sex-biased migration due to female neoteny

We investigated sex-biased migration by searching for mitochondrial to nuclear phylogenetic incongruency. The maternal (mitochondrial) relationships between populations revealed one fundamental incongruence with the nuclear tree: the Swiss population being sister to the Germany (**Figure 4B**), whereas in the nuclear tree the Italian population is sister to the German populations (**Figure 4A**). We further investigated sex-biased migration by estimating PoMo population trees from X-linked contigs. The X-linked contigs corroborated the mitochondrial history, where the Swiss population is sister to the Germany in 57% of the contigs. The Italian population was sister to the Germany in 43% of the X-linked contigs (**Figure 4C, Figure S8**). In contrast, the autosomal contigs showed a strong Italian-German sister relationship (89% of contigs). The uncovered nuclear-mito-X incongruency proposes at a scenario of sex-biased migration.

### Testing for population gene-flow

Our STRUCTURE/ADMIXTURE analyses showed little evidence of gene flow (**Figure 2B, Figure S1, S2**), which is contrary to the expectation of one single widespread European species with recent, post-glacial expansion. On the other hand, strong population structure and low gene-flow matches the expectation of limited migration due to female neoteny. We performed a *f-branch* statistic to assess the proportion of shared alleles across populations. We tested migration events assuming the most probable autosomal population tree, where the Italian population is sister to the German populations, and the most probable mitochondrial population tree, where the Swiss population is sister to the German populations. In the first scenario, we identified two putative migration events, the first between the ancestral population of Italy and Germany with the Finish population, sharing 8% of alleles, and the same ancestral population with the Swiss population, sharing 14% of shared alleles (**Figure 5A**). In the second scenario, we identified multiple migration events. First, three migration events between the Finish population and ancestral population of the Jena and Kiel populations (<1% of shared alleles), the ancestral populations of all German populations (2% of shared alleles), and the ancestral population of the Swiss and German populations (7% of shared alleles). Second, a migration event between the Swiss population and ancestral population of the Jena and Kiel populations (1% of shared alleles). Third, a migration event between the Swiss population and the Italian populations (1-2% of shared alleles). The *f*-branch statistics revels different patterns of shared alleles in the autosomes and mitochondria.

### ABC analysis

We used an ABC framework to explore the contribution of a ghost population as a founder ancestral population and to further assess the population relationships between Germany – Italy – Switzerland. We tested the following models: *Model 1*: Finland as the most ancestral population from which Switzerland and Italy are derived independently. Switzerland is sister to Germany. *Model 2*: Finland as the most ancestral population from which Italy derives. Switzerland and Germany derive from the Italian population. Italy is sister to Germany. *Model 3*: Finland as the most ancestral population from which Switzerland derives. Italy and Germany derive from the Swiss population. Switzerland is sister to Germany. *Model 4*: A ghost population represents the most ancestral population, from which Finland and Switzerland derive independently. Italy and Germany derive from the Swiss population and Switzerland is sister to Germany.

Using 12,572 SNPs retrieved from intergenic regions, we calculated the joint site-frequency spectrum (JSFS), which showed that there is little to no fixed differences between pairs of populations (especially for populations pairs that exclude Finland), and that most of the polymorphisms present in this dataset are either private or shared between each pair of populations (**Table S2, S3**). Posterior probabilities for the four tested models were the following: *Model 1* (0.000), *Model 2* (0.000), *Model 3* (0.016), and *Model 4* (0.984). Parameter estimation on *Model 4* showed that the effective population size is around 1.448 million individuals and that the German population split from the Swiss about 266k generations ago.

Moreover, the Italian population split from a putative ghost population about 944k generations ago. On the other hand, the Swiss population split from the Italian population about 446k generations ago, and the Finish population split more recently from the same ghost population about 614k generations ago (**Table 2**). ABC was not able to retrieve accurate estimates of population-specific *N*_e_’s.

**Table 2.** ABC parameter estimates and their respective priors. These posterior distributions are shown in **Figure S9**. Here, N_e_ stands for effective population size, *T* corresponds to the split time of each population.

| Parameter | Prior | Mode | 95% quantiles |
| --- | --- | --- | --- |
| $\log_{10}(N_e)$ | unif(4.5,7.5) | 6.161 ($N_e=1,448,772$ ind.) | (4.43,7.01) |
| $T_{\text{GeGl}}$ | unif($0.1*20000/4N_e$, $10*20000/4N_e$) | $0.046*4N_e$ generations ago | (0,1.42) |
| $T_{\text{SwLa}}$ | unif($T_{\text{GeGl}}$, $10*25000/4N_e$) | $0.163*4N_e$ generations ago | (0,3.47) |
| $T_{\text{ItCo}}$ | unif($T_{\text{SwLa}}$, $10*40000/4N_e$) | $0.077*4N_e$ generations ago | (0,2.20) |
| $T_{\text{FiHe}}$ | unif($0.1*40000/4N_e$, $10*40000/4N_e$) | $0.106*4N_e$ generations ago | (0,2.23) |

## Discussion

The big European firefly *L. noctiluca* is a popular insect species, widely distributed across Europe, where neotenic females signal their presence to males by producing light. Female neoteny limits migration and gene-flow to one sex, as only winged males can disperse at the adult stage. We used a genomic approach to investigate population structure, past demographic events and gene flow in *L. noctiluca* with the main goal of understanding the impact of sex-biased migration into present nucleotide diversity patterns in insects.

Our results inferred variable effective population sizes over the last 100k years ranging from a maximum of 6.3 million individuals to a minimum of 7k individuals most likely severly affected by the last ice age (**Figure 3A**). Unsurprisingly, the large effective population sizes lead to old estimates of the common ancestor of individuals from the same population (∼800 kya). The common ancestor of individuals from the same population was in fact the same as between population, as shown by the strong correlation of estimates per contig compared with between contigs (**Figure 3B**). Thus, populations of *L. noctiluca* share ancestral polymorphisms and the divergence between populations must have occurred after (i.e., more recent) than the age of the common ancestor. Despite the presence of ancestral polymorphisms, we inferred high levels of population structure with little gene flow using conventional approaches (STRUCTURE, ADMIXTURE, TreeMix and *f*-branch statistic). Our new polymorphism aware phylogenetic divergence time estimation (PoMos) using full genome and all samples per population, inferred the main population-splitting events in the last ∼40k years, thus during the last ice age instead of after it. Our most exciting result is the nuclear-mitochondrial-X incongruency of the population history. The mitochondrial and X-linked analyses of all samples per population suggest an Western European colonialization of Germany possibly from a French population. Only later, likely after the retreat of the glaciers in the European Alps, a secondary contact between the Italian and German populations was established. Interestingly, we uncovered evidence of a male-only migration, as not a single mitochondrial genome from a German population suggested Italian ancestry (**Figure 4B**).

### High levels of population structure and species status

We uncovered high levels of population structure where populations showed strong differentiation according to country. Only the three German populations formed a single genetic cluster, showing the lowest retrieved *F*_ST_ values (0.08 – 0.11) (**Figure 2**). The German populations presented recent divergence times, which suggest that Germany was colonized by a single event. We suggest a south to north migration pattern for the German populations, reflected by the correlated pattern of shared alleles to latitude (ADMIXTURE, k4-k5) and by the binomial distribution of the permutation Mantel test (**Figure Sx, Sx**). The remaining populations formed defined genetic clusters with high *F*_ST_ values ranging 0.21 – 0.48, suggesting low levels of gene flow. Similarly high *F*_ST_ values have been retrieved for between species comparisons, such as between different species of *Heliconius* butterflies (Van Belleghem et al. 2018) or between closely related fox species (L. Rocha et al. 2023). This observation raises the question of the speciation status across populations of *L. noctiluca* and the effect that neoteny can have in the process of speciation.

### Population history and and the addition of a ghost population

The estimated PoMos tree (**Figure 4**) placed the Finish as the population with the deepest divergence, followed by Switzerland, Italy and Germany, a topology that rules out an isolation by distance, stepwise colonization pattern from south to north. The deep divergence of the Finish population and our ABC analysis suggests that the founder of this populations is only distantly related to the rest of the sampled populations. According to *L. noctiluca’s* geographical distribution, the tested ghost population (*Model4*) could correspond to an Eastern population, colonizing Finland (**Figure 5B**). Interestingly, the addition of an Eastern population as a proxy for a glacial refugia population resulted in the Italian population showing an older divergence time than the Finish population. The divergence time of the Swiss population was about ∼25kya, which corresponds to a period when Switzerland was covered by a glaciar (Seguinot et al. 2018), surely hindering colonization at that time. The relatively deep divergence time of the Swiss population poses the hypothesis that a Western population might be the founder of Switzerland. Sampling *L. noctiluca* populations from Western and Eastern European regions will further shed light on the population history and dynimics of this species. The above presented divergence times should be treated cautiously. The mutation rate is not known for this species yet, therefore the estimated divergence times can change accordingly. Additionally, the estimated divergence times also depend on the model of population size changes used and the force of selection. Nevertheless, the present study constitutes the first to estimate divergence times using full genomes and multiple-population data and the first to propose a comprehensive demographic species for *L. noctiluca*.

### Shared alleles across populations

Using an *f-*branch statistics approach, we were able to detect putative migration within the Italian-German internal branch and two populations: Finland and Switzerland, reaching 8 and 14% of migration fraction, respectively. The detection of migration involving an internal branch can be interpreted as migration events happening in the past or between unsampled lineages (Malinsky et al. 2018; Suvorov et al. 2022). We consistently found migration between an internal branch and Finland, as further explored with an *f*-branch analysis based on the mitochondrial tree (**Figure 5A**) and by a TreeMix analysis (**Figure S10**), further suggesting a connection with an unsampled population or past migration with Finland. Our population structure and gene-flow analysis suggest high population structure between the collected populations with the presence of putative ancestral gene-flow. Currently, we do not have information about the generational displacement of larvae or adults, information that would contribute to the comprehension of the migration biology of *L. noctiluca*. Sampling of additional populations will contribute to have more information on migration breadth of this species.

### Sex-biased migration

We uncovered a nuclear-mitochondrial-X chromosome tree discordance, where the sister population to Germany differs according to the genomic region tested. The mitochondrial tree shows a scenario where Germany was initially colonialized by the ancestral population of the Swiss population, possibly of Western European origin. In such a scenario, female larva are the ones that set the limit for a range expansion. The autosomal signal on the other hand, suggests a scenario where Italy is closer related to Germany, a signal which we hypothesize might be driven by male-biased migration between Italy and Germany. The X chromosome nicely captures both, the autosomal (43%) and mitochondrial (57%) signals, further supporting a male-biased migration scenario. The divergence times of the X-linked contigs supporting a Swiss-German split is older than that of the X-linked contigs supporting the Italian-German split, further supporting a putative German colonization by Western European population. The detection of nuclear-mitochondrial-X chromosome incongruence shows the putative effect that neoteny can have on colonization and migration patterns. Currently, we do not have information about the generational displacement of larvae or adults, information that would contribute to the comprehension of the migration biology of *L. noctiluca*. The sampling of additional populations connecting our populations sampled in this study have the potential of revealing the migration breadth in this species.

## Conclusions

Female neoteny, which limits dispersal capability, can play an important role in a species’ range expansion and migration. In this work, we studied the demographic history and sex-biased migration of the big European firefly *L. noctiluca*. We generated a high-quality genome assembly, which includes PacBio IsoSeq gene annotation, transposable elements characterization and the identification of the X chromosome. Population sampling of 115 individuals from Italy to Finland produced the first results on population structure, nucleotide diversity levels, migration rates and demographic inference of *L. noctiluca*. We found very strong population structure with very low migration between populations. We applied a novel approach for full-genome population data, polymorphism aware phylogenetic models (PoMo), to estimate divergence times, population relationships and discordance among genomic regions. Our results demonstrate that range expansion is followed by migration from males, the sex with higher dispersal capability (introgression). The migration signal was not detected by traditional approaches and sex-biased migration should be considered for future model and hypothesis development. Finally, the population colonization of *L. noctiluca* is more complex than expected, where we hypothesize that unsampled populations from western, eastern and southern Europe have a big potential of further explaining the colonization history of this species.

## Methods

### Sample collection, DNA extraction and sequencing

We sampled specimens in seven locations in Europe. Males were collected using a funnel light trap with a yellow led lamp (2.0-2.2V). Light traps were put in the ground short after sunset and left on for two hours. Females were collected by walking along transects. Collected individuals were stored in 96% ethanol at -4°C. For each population 15-20 individuals were collected. The single Greek individual from an unknown *Lampyris* species served as an outgroup.

High molecular DNA was extracted using the kit MagAttract HMW DNA kit (Qiagen) following manufacturer’s guidelines. DNA fragment sizes and integrity were checked with a 1% agarose gel and a Femto Pulse system (Agilent). Long DNA fragments were sequenced from a single male individual using Nanopore PromethION in one flongle cell and run for 72 hours. Nanopore sequencing was done by the SciLifeLab in Uppasala, Sweden.

Short molecular DNA was extracted using the Monarch Genomic DNA Purification kit (New England BioLabs). DNA quality and integrity were assessed with a Nanodrop and an Agilent 5400. Illumina 150bp paired-end reads were generated for each sample, aiming at a 15xcoverage, with the exception of the genome’s individual from which 60x sequencing depth was generated. Library prep and sequencing was outsourced to Novogene, China.

### De novo genome assembly

Base calling for Nanopore reads was done with Guppy (4.0.11). A hybrid genome assembly was performed with MaSuRCA v4.0.5 (Zimin et al. 2013) using 15Gb of Nanopore reads and 60x depth 150bp paired-end Illumina reads from the same individual, followed by two rounds of haplotype purging with Purge_dups v1.2.5 (Guan et al. 2020). Sequences not belonging to the class Insecta were identified using Blobtools v1.1.1 (Laetsch et al. 2017) and removed from the assembly. No genome polishing was performed for *L. noctiluca*, as further polishing lead to worse BUSCO scores. Genome size and genome heterozygosity levels were estimated with GenomeScope (Vurture et al. 2017). Genome statistics were calculated with Quast v5.0.2 (Gurevich et al. 2013) and genome completeness was assessed with BUSCO v5.2.2 (Simão et al. 2015) using the dataset Insecta. Repeats were annotated using RepeatModeler v1.0.11 and the generated custom made repeat library was used for genome masking with RepeatMasker v 4.1.2 (Smit et al. 2015).

### Identification of the X chromosome

To identify the putative contigs belonging to the X chromosomes we compared male to female (m:f) coverage ratio across contigs. *L. noctiluca* males are expected to be the heterogametic sex, thus we expect a m:f coverage ratio on the X to lie near 0.5. We used Illumina reads from 2 samples of each sex. Read quality control was done with FastQC (V 0.11.9) and trimming of adaptors and tails was done with Cutadapt (V 3.4) using a threshold of Phred < 20. The curated reads were mapped to the hard masked genome (Repeat Masker, V4.1.2) with BWA (V0.7.17). Duplicate reads were removed from the bam files with Picard (V 2.20.8). Coverage was calculated with Deeptools (Version 3.5.0) for 10Kb windows across the genome and normalized using RPKM (Reads Per Kilobase Million). Each 10kb window coverage level was normalized by dividing it by the mean coverage value of the 5 five largest autosomal contigs. These five contigs were manually selected by choosing the five largest contigs with a male to female coverage ratio of 1 ± 0.1. Contigs smaller than 30kb were filtered out leaving only contigs with at least 3 data points (i.e three 10kb windows). We then performed a non-parametric Wilcoxon Rank Sum Test to test for significant differences in contig coverage values between sexes and applied a Bonferroni multiple test correction. Male to female coverage ratios were calculated only from contigs with significant differences in coverage between sexes. Contigs with a m:f ratio 0.4 ≤ x ≥ 0.6 were considered to belong to the X chromosome.

### Identification of coding sequences using PacBio IsoSeq transcriptome data

RNA was extracted from heads and thorax+abdomen of one female and one male using the Monarch Total RNA Miniprep Kit (New England, BioLabs). RNA was purified by ethanol precipitation and equal concentration of head and thorax+abdomen tissue was pooled for sequencing, separately for each sex. Equal concentration of each body type ensures equal probability of transcript sequencing. IsoSeq libraries and sequencing were done by Novogene where 69 and 91 subread bases (Gb) were produced for the male and female sample, respectively. Primer removal, multiplexing of raw reads and clustering was done with IsoSeqv3 v3.8.0 (https://github.com/PacificBiosciences/IsoSeq). In order to have a single non-redundant transcript set for the species, transcript collapsing was performed with Cupcake v29.0.0 (https://github.com/Magdoll/cDNA_Cupcake) and BUSCO in transcriptome mode was run to assess transcriptome completeness. Curated transcripts were mapped back to the genome with minimap2 v2.14 (Li 2018) and a gft file was produced denoting intergenic, exons and intronic regions (**Appendix 1**).

### Processing of population level whole genome re-sequencing

Illumina short read sequences were trimmed with TrimmGalore! V0.6.6 (Krueger 2012) and FastQC v0.11.9 was used to filter out bases with a phred score < 20 and reads shorter than 20bp (Andrews 2010). Reads were mapped back to the genome with BWA v0.7.17 (Li and Durbin 2009). Mapped files in BAM format were curated by removing PCR duplicates with Picard v2.20.8, and low-quality reads (Q20) were discarded using SAMtools v1.10 (Li *et al*. 2009).

GATK v4.1.9 (Auwera et al. 2013) was used to call SNPs and indels via local re-assembly of haplotypes with HaplotypeCaller. Joint genotyping of all sequenced samples was done with GenotypeGVCFs. VCFs statistics were drawn with bcftools stats and gatk VariantsToTable. Quality scores thresholds were applied for minimum and maximum read depth [20,1568], fisher strand [FS=10], strand bias [SOR=3], root mean square mapping quality [MQ=40] and nucleotide quality by depth [DP=2]. Only variants with a QUAL > 30 were kept, as well as only SNPs (indels were removed) and biallelic sites. A SNP missingness of 0.25 across all samples was allowed. Sites in the VCF file which overlapped with repetitive elements were excluded from the analysis. An additional set of VCF files which included monomorphic sites was generated, where the GATK tag --select-type-to-include NO_VARIATION was used.

### Population genetic and structure analysis

Population genetic analysis were done separately on the autosomes and the X chromosome. Nucleotide polymorphism diversity (depicted by π per site) and Tajima’s D were estimated with VCFtools v0.1.14 (Danecek et al. 2011), separately for exons, introns and intergenic regions on sliding windows of 10000 base pairs.

Population structure was first explored via Principal Component Analysis (PCA) in Plink v1.09 (Chang et al. 2015), filtering out linked sites with r^2^ > 0.2. ADMIXTURE (Alexander et al. 2009) and fineSTRUCTURE (Lawson et al. 2012) were run using the unlinked SNP set, for K1-K6. Population differentiation was calculated as Fst (Weir and Cockerham 1984) across all population pairs with VCFtools in windows of 10000bp. Genetic distances (Dxy) (Wakeley 2016) were calculated for every pair of populations using pixy (Korunes and Samuk 2021).

### Population size estimation

We estimated population size trajectories using the StairwayPlot approach (Liu and Fu 2020) within a Bayesian statistical framework as implemented in RevBayes. We used SNPs from the autosomes as data, comprising a total of 56-59 million SNPs per population. We assumed a total genome size of 411.5Mb for the computation of monomorphic sites. This number of monomorphic sites is lower than the total genome size as we estimated that 30-40% of the variable sites were filtered out, thus we reduced the corresponding genome size accordingly. The number of variable sites is informative about the actual effective population size but not about the population size changes over time. We explored the impact of data filtering and different genomic regions by performing the StairwayPlot analysis for all SNPs (main results), and only intergenetic regions, exons or introns.

For each population we computed the site frequency spectrum in RevBayes directly from the VCF file. The complete site frequency spectrum had between 30 (for Finland) to 40 (Italy and Germany) categories. We used the folded site frequency spectrum because the sites were not polarized.

We assumed a mutation rate of 2.8E-9 per site per year and a generation time of one year. Our prior model for the population size trajectory assumed autocorrelated, log-normal distributed displacements with an expectation of one order of magnitude over the total timespan. We explored several additional prior models, including uncorrelated models where population sizes per interval are drawn from a lognormal prior distribution. We ran a Markov chain Monte Carlo simulation for 250,000 iterations with sampling every 10 iterations. We applied the same settings for all populations. We plotted the resulting population size trajectories using the R package *RevGadgets*.

### Coalescent estimations of the last common ancestor

We estimated the time of the most recent common ancestor of all sampled individuals within a population using a genealogy-based coalescent approach as implemented in RevBayes. We constructed multiple sequence alignments including invariant sites per contig for each population. We inferred the genealogy based on the alignments using the following model assumptions. We assumed a standard phylogenetic GTR+I nucleotide mutation model. We assumed a coalescent process prior on the genealogy and a “known” mutation rate of 2.8E-9 per site per year. We ran four replicated Markov chain Monte Carlo analyses for 50,000 iterations each with 197 moves per iteration. We checked for convergence using the R package *convenience*. From the posterior sample of genealogies, we extracted the root age to represent the time of the most recent common ancestor. We performed this genealogy-based coalescent analysis for 33 contigs from the autosomes and 34 contigs from the X chromosome.

### Demographic inference with polymorphism aware phylogenetic models

We estimated the population relationship and divergence times between populations using polymorphism aware phylogenetic models (PoMo) implemented in RevBayes. PoMo models use as data allele counts per population and therefore can handle multiple individuals per population efficiently. Changes in allele frequencies are modelled using a Moran process combined with a boundary mutation process. Instead of a 4-state nucleotide process we converted the data into binary states. We used a total of 61,441,738 SNPs from the autosomes and implemented an ascertainment bias correction for not using monomorphic sites. We assumed a “known” mutation rate of 2.8E-9 and an average effective population size of 100k. In further analyses we explored the impact of the *a priori* assumed average effective population size. We assumed a uniform prior distribution on both topology and divergence times. We ran two replicated Markov chain Monte Carlo analyses for 50,000 iterations each with n moves per iteration. We computed the *maximum a posteriori* topology and mean divergence times from the posterior samples. We checked for convergence using the R package *convenience*. We plotted the population tree using the R package *RevGadgets*.

### Demographic inference with Approximate Bayesian Computation (ABC)

#### Data collection

Data consists of single nucleotide polymorphisms (SNPs) obtained from 10 randomly chosen intergenic regions coming from four populations of *L. noctiluca*. The populations are Helsinki-Finland (FiHe), Lausanne-Switzerland (SwLa), Italy (ItCo), and Germany (GeGl). A total of 12572 SNPs coming from the intergenic regions were kept for downstream analyses (**Table S4**). The “FiHe” population consisted of n=15 individuals, the “SwLa” population with n=20 individuals, the “ItCo” population with n=20 individuals, and the “GeGl” with n=19 individuals, yielding a total of 74 sampled individuals. Recombination rates for each of the intergenic regions were estimated using the software *LDhat* (Auton and McVean 2007).

#### Observed summary statistics

We calculated a total of 370 summary statistics, including: number of segregating sites *S*, Watterson’s *θ*_W_ (Watterson 1975), *π*, Tajima’s *D* (Tajima 1989b), linkage disequilibrium *Z*_nS_ (Kelly 1997), the folded SFS, Weir-Cockerham’s *Fst* (Weir and Cockerham 1984), distance of Nei (Nei and Li, 1979), and the Wakeley-Hey “W” summaries of the joint SFS (Wakeley and Hey 1997). All the above-mentioned statistics are unaffected by the polarization (or lack thereof) of the observed SNPs.

#### Demographic models

We tested four different demographic scenarios: scenario 1) FiHe and SwLa split from an ancestral Finish population, then ItCo splits directly from FiHe, and finally GeGl splits from SwLa; scenario 2) FiHe and ItCo split from an ancestral Finish population, then SwLa splits from ItCo, and finally GeGl splits from ItCo; scenario 3) same as scenario 2, but GeGl splits from SwLa 4) Same as scenario 3, but both FiHe and ItCo split from a putative “Eastern” population (Figure 1). With these different demographic scenarios, we covered biologically plausible population histories of European *L. noctiluca*.

#### ABC simulations

We performed simulations with the program *msms* (Ewing and Hermisson 2010). For each of the four demographic models described above (Fig. 1) we simulated segregating sites for 74 individuals (15, 20, 20, and 19 individuals representing the “FiHe”, “SwLa”, “ItCo” and “GeGl” populations, respectively). From the simulated sites we calculated all summary statistics described above. All priors are shown in Table 4. We repeated this whole simulation process 20,000 times.

#### Model choice and parameter estimation

With all 20,000 simulations per model we calculated the posterior probabilities of each of the four demographic scenarios using the R package *abc* (Csilléry et al. 2012). Model choice was based on the following summary statistics per population: *θ*_*W*_, *π*, Tajima’s *D, Z*_*nS*_, W statistics, distance of Nei, and *Fst*. Parameter estimation on the best model was accomplished by using both the rejection (Tavare et al. 1997) (Pritchard et al. 1999) and regression (Beaumont et al. 2002) algorithms using the same R package *abc*. To reduce dimensionality while keeping the maximum amount of information still available we used partial least squares (pls) in the context of ABC (Wegmann et al. 2009).

### Mitochondrial genome tree inference

We constructed a multiple sequence alignment for the mitochondrial genome comprising of 115 sequences of 18,937bp each. We estimated the mitochondrial relationship using a standard phylogenetic approach as implemented in RevBayes. We assumed a GTR+GI mutation model with 4 rate categories (slow to faster evolving sites). We assumed a mitochondrial mutation rate of 1.34E-8 per site per year. We assumed a uniform prior distribution on the root age, the topology and node ages between lineages. We ran two replicated Markov chain Monte Carlo simulations for 50,000 iterations with 614.6 moves per iteration, sampling every 10^th^ iteration. We checked for convergence using the R package *convenience*. We plotted the mitochondrial tree using the R package *RevGadgets*.

### Analysis of gene-flow

We performed several types of analysis to infer gene-flow and migration between the sampled *L. noctiluca* populations. Our primary analysis consisted of a polymorphism aware phylogenetic model (PoMo) analysis, which we performed separately for 35 autosomal and 34 X-linked contigs. These contigs were chosen based on length. For each contig, we computed for each SNP the allele frequency per population and used the allele frequencies as data. We used an ascertainment bias correction to condition on only variable sites. We ran two replicated Markov chain Monte Carlo analyses for 50,000 iterations. We computed the posterior probabilities of the Italian and the Swiss population being sister to the German populations.

Additionally, we explored putative migration events between the populations using the *f4*-branch statistic as implemented in Dsuite (Malinsky et al. 2021). As input data, we used the previously curated VCF file including monomorphic sites. To disentangle correlated *f4*-ratio results and to assign evidence of gene flow across the phylogenetic tree we used the *f*-branch metric implemented by the same authors. The most probable phylogenetic tree was used for the analysis, where Italy is sister group to Germany. A Treemix (Pickrell and Pritchard 2012) analysis was done for autosomal unlinked SNPs using the following command: treemix –I $TreeInput -m $i -o lanoc_M_.${i} -root GrPa -bootstrap -k 1000. Treemix was run to investigate up to six migration events (six edges) using the Greek individual to root the population tree and allowing for 1000 replications.

## Acknowledgments

We would like to thank Gabriele Kumpfmüller for her magnificent technical assistance in the lab. Andreas Tiraboschi (Gruppo Ecologico Colognese) for sharing his knowledge with us on Italian fireflies. Klaus Reinhold for guiding and showing us where to find fireflies in Greece. To Shanaka Thisara for his fine imaging skills of *L. noctiluca* specimens. This work was funded by the Société Vaudoise des Sciences Naturelles (SVSN) and the Société Académique Vaudoise (SAV) to PD and by the DFG SPP-1991 to SH and AC.

