## Supplementary material for "Sex-biased migration and demographic history of the big European firefly *Lampyris noctiluca*": Suplementary file

*Supplementary Tables* *pages 2-3*

*Supplementary Figures.* *pages 4-10*

### Supplementary tables

**Table S1.** Genome statistics of *Lampyris noctiluca*'s genome assembly using different assembly approaches.

| Assembler | MaSuRCA | MaSuRCA after curation | Flye purged | Canu | Shasta |
|---|---|---|---|---|---|
| Genome size (bp) | 658,164,932 | 654,253,030 | 757,198,623 | 717,629,941 | 559,715,407 |
| No. of contigs | 1500 | 1416 | 9033 | 3597 | 6739 |
| N50 | 649,158 | 653,271 | 287,897 | 439,832 | 217,701 |
| L50 | 383 | 293 | 669 | 468 | 737 |
| Busco_insecta | 94.60% | 94,70% | 90.70% | 70.20% | 49.20% |

**Table S2.** Mean observed summary statistics over 10 intergenic regions (Table 1). Sample sizes per population are given in parentheses.

| | **FiHe (n=15)** | **SwLa (n=20)** | **ItCo (n=20)** | **GeGl (n=19)** |
|---|---|---|---|---|
| ***S*** | 332.1 | 256.5 | 358.1 | 391 |
| $\theta_W$ | 102.14 | 73.39 | 100.93 | 110.21 |
| $\pi$ | 95.74 | 65.15 | 81.96 | 87.20 |
| **Tajima's *D*** | 0.0056 | -0.58 | -0.62 | -0.88 |
| $Z_{nS}$ | 0.149 | 0.188 | 0.078 | 0.134 |

**Table S3.** Population pairwise statistics. Wakeley-Hey *W* statistics summarizing the Joint-SFS, plus population differentiation statistics.

| | **FiHe-GeGl** | **FiHe-ItCo** | **FiHe-SwLa** | **GeGl-ItCo** | **GeGl-SwLa** | **ItCo-SwLa** |
|---|---|---|---|---|---|---|
| W1 (polymorphic in population 1) | 261.4 | 266.1 | 285.9 | 404.4 | 352.8 | 528.2 |
| W2 (polymorphic in population 2) | 346.6 | 318.4 | 236.6 | 65 | 42.4 | 26.4 |
| W3 (fixed differences in both populations) | 14.9 | 14.6 | 36 | 0 | 0 | 0 |
| W4 (shared polymorphisms in both populations) | 253.2 | 248.3 | 228.7 | 616.8 | 633.9 | 1222.4 |
| $F_{st}$ (Weir and Cockerham) | 0.187 | 0.194 | 0.282 | 0.115 | 0.107 | 0.05 |

**Table S4.** Selected intergenic regions for the ABC analysis, and number of SNPs for all populations combined.

| **Intergenic region** | **From** | **To** | **#SNPs** |
|---|---|---|---|

| | | | |
|---|---|---|---|
| scf7180000016868_F_17831_scf7180000022641_F | 1807660 | 2108297 | 3809 |
| scf7180000017133 | 641048 | 736029 | 1592 |
| scf7180000017976 | 1 | 569585 | 4630 |
| scf7180000019093 | 1 | 259092 | 2104 |
| scf7180000019103_F_2937_scf7180000017693_F | 555401 | 582998 | 53 |
| scf7180000020236 | 267456 | 283519 | 175 |
| scf7180000020969_F_8272_scf7180000020710_F | 409437 | 425440 | 82 |
| scf7180000021855_F_1147_scf7180000021523_F | 82952 | 84155 | 11 |
| scf7180000022443_R__3794_scf7180000017785_F | 514446 | 539765 | 14 |
| scf7180000022736_F_2769_scf7180000023558_R | 157979 | 170766 | 102 |
| **Total SNPs** | - | - | **12572** |

### Supplementary figures

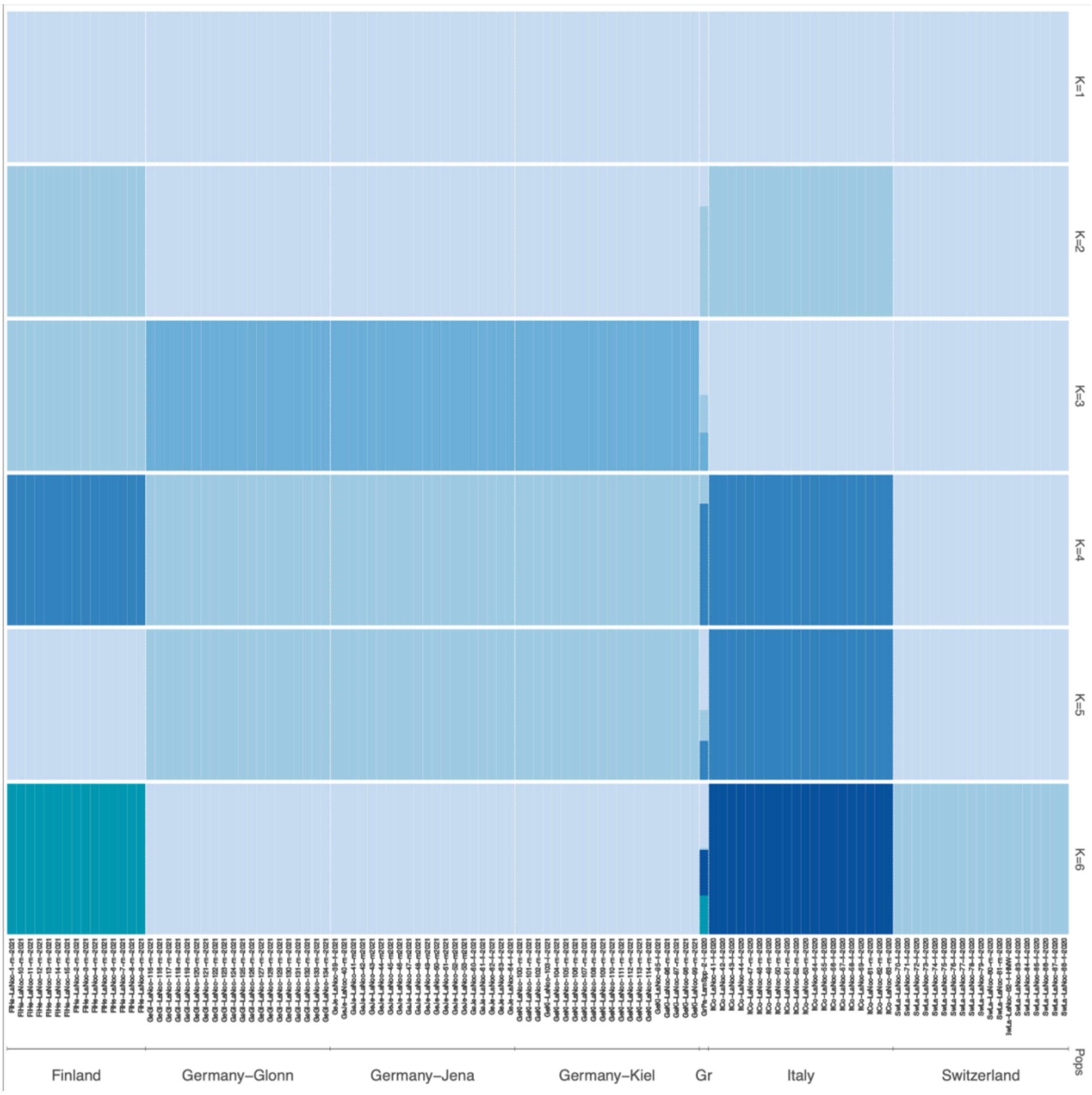


**Figure S1.** STRUCTURE analysis showing genetic clustering of the autosomes conditioning for k1-6 of six populations of *L. noctiluca* and an outgroup individual.

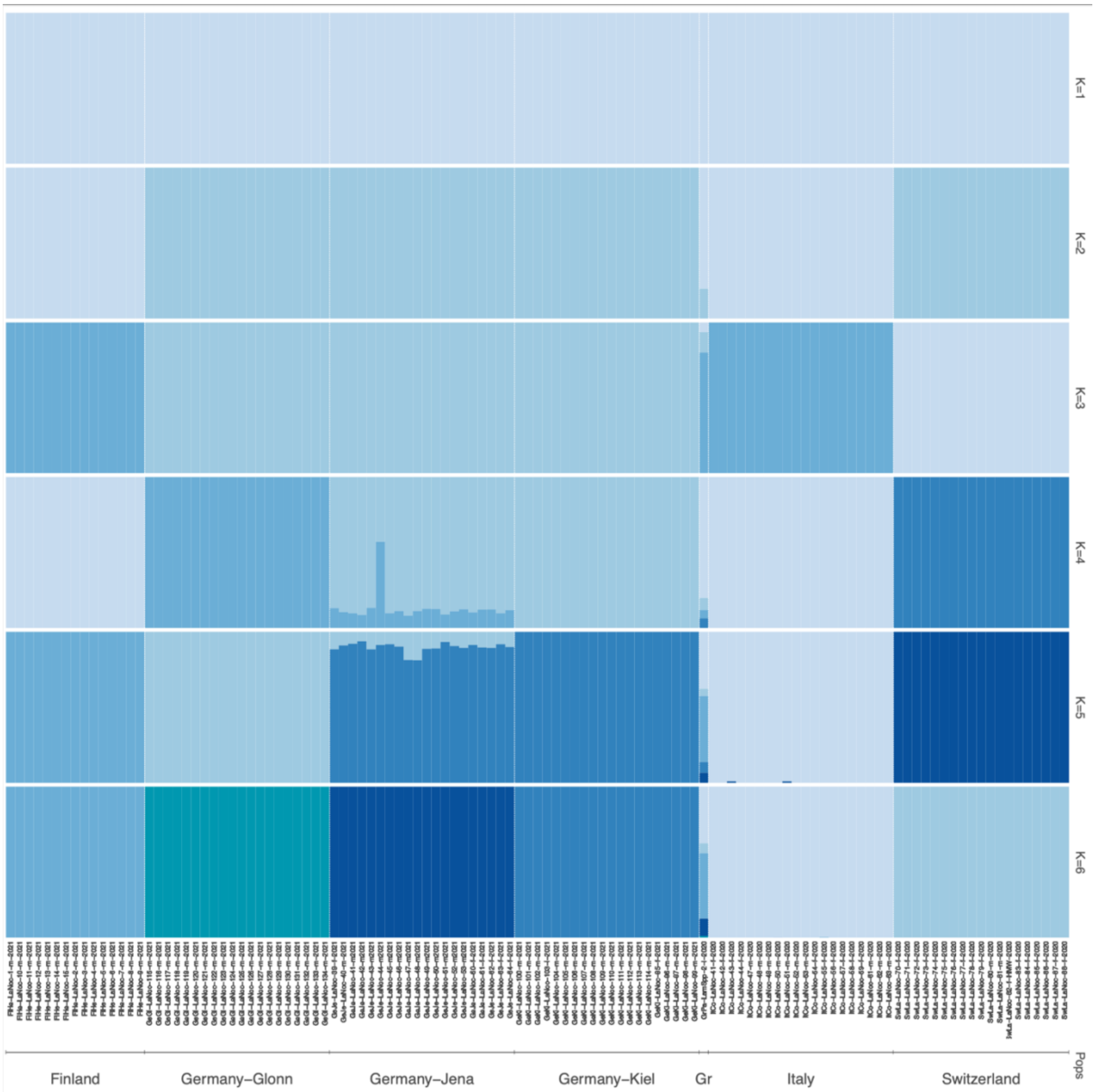


**Figure S2.** ADMIXTURE analysis showing genetic clustering of the autosomes conditioning for k1-6 of six populations of *L. noctiluca* and an outgroup individual.

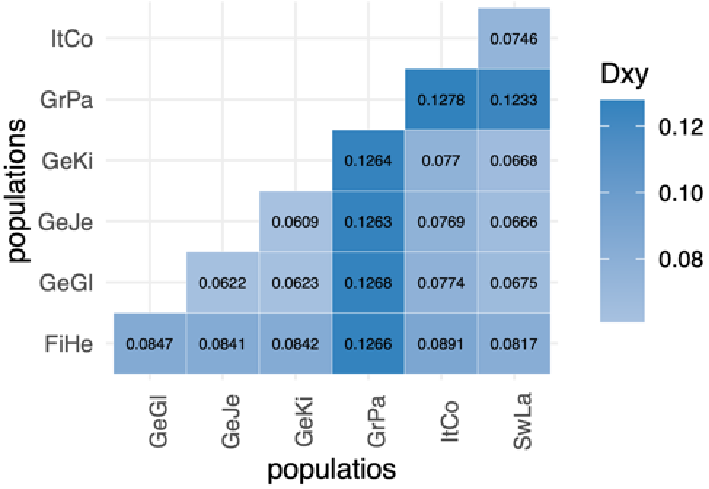


**Figure S3.** Nei's absolute genetic divergence (*Dxy*) for every population pairwise comparison. Color gradient represents the level of genetic divergence. FiHe: Finland, GeKi: Germany-Kiel, GeJe: Germany-Jena, GeGl: Germany-Glonn, ItCo:Italy, SwLa: Switzerland.

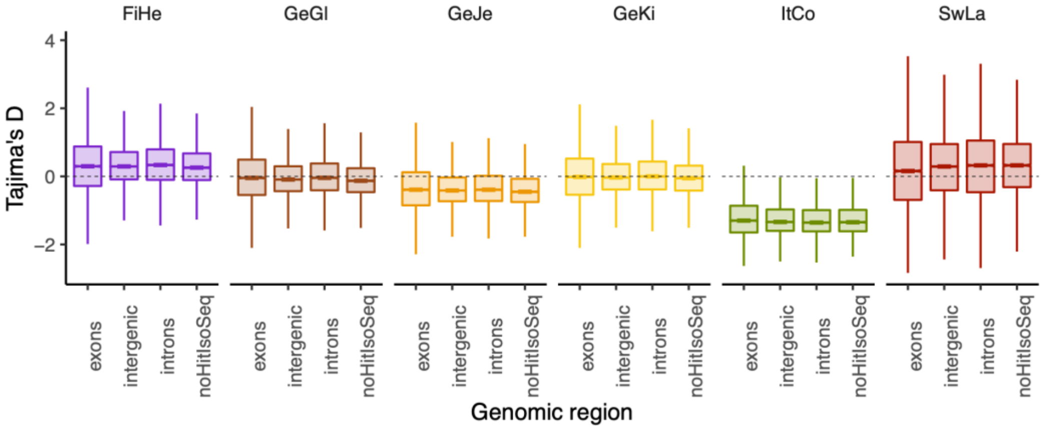


**Figure S4.** Boxplots showing the distribution of Tajima's D for the autosomes (upper panel) and the X chromosome (lower panel). Tajima's D was estimated for different genomic regions. Contigs labeled as noHitIsoSeq, are contig were no hits IsoSeq  transcripts were found. Most probably contigs with no IsoSeq hits belong to the intergenic region category.

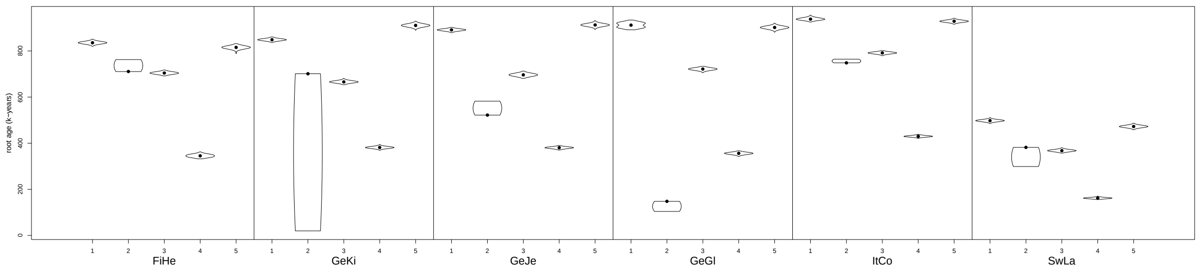

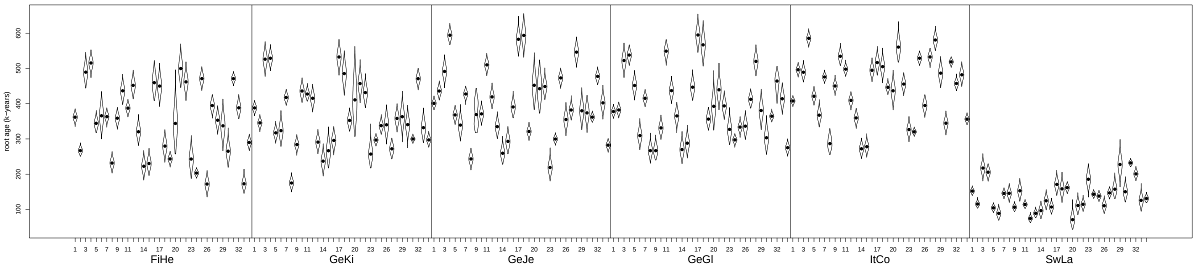


**Figure S5.** Coalescent ages for the autosomes (upper panel) and the X chromosome (lower panel)

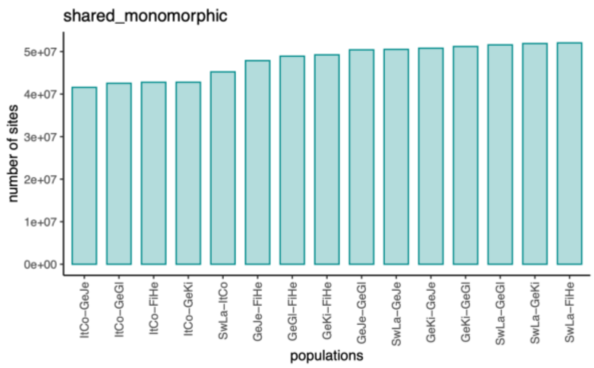


**Figure S6.** Pairwise estimation of shared polymorphisms for each tested population.

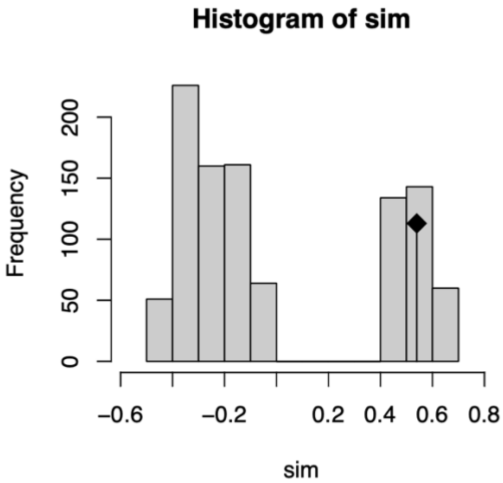


**Figure S7.** Histogram showing the frequency of permutations between population genetic distances as measured by *Fst* and geographic distances. The observed value is shown as a tilted black square. Mantel test *p*-value= 0.137.

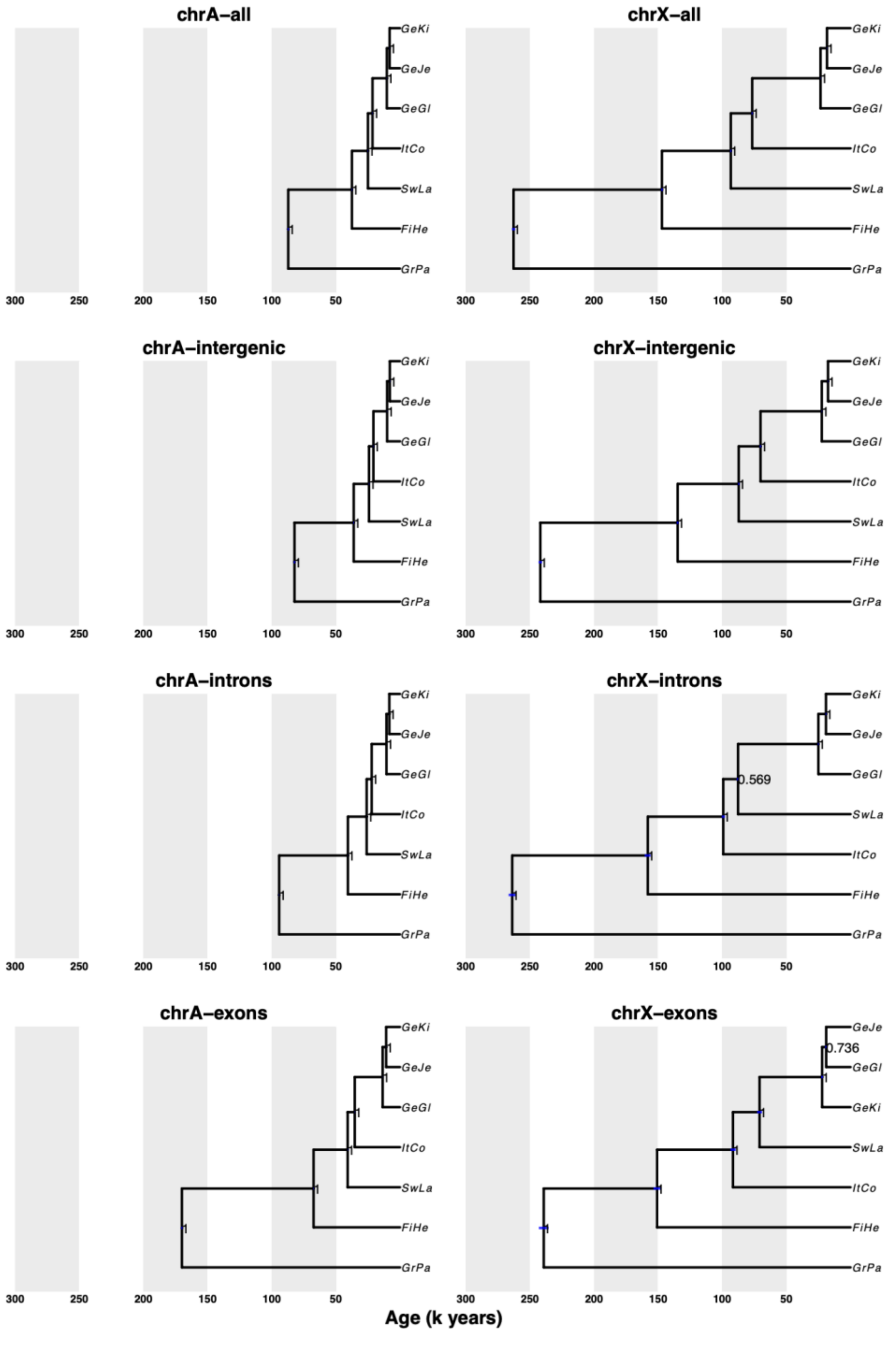


**Figure S8.** PoMo analysis of the autosomes (chrA) and the X chromosome (chrX) using different genomic regions.

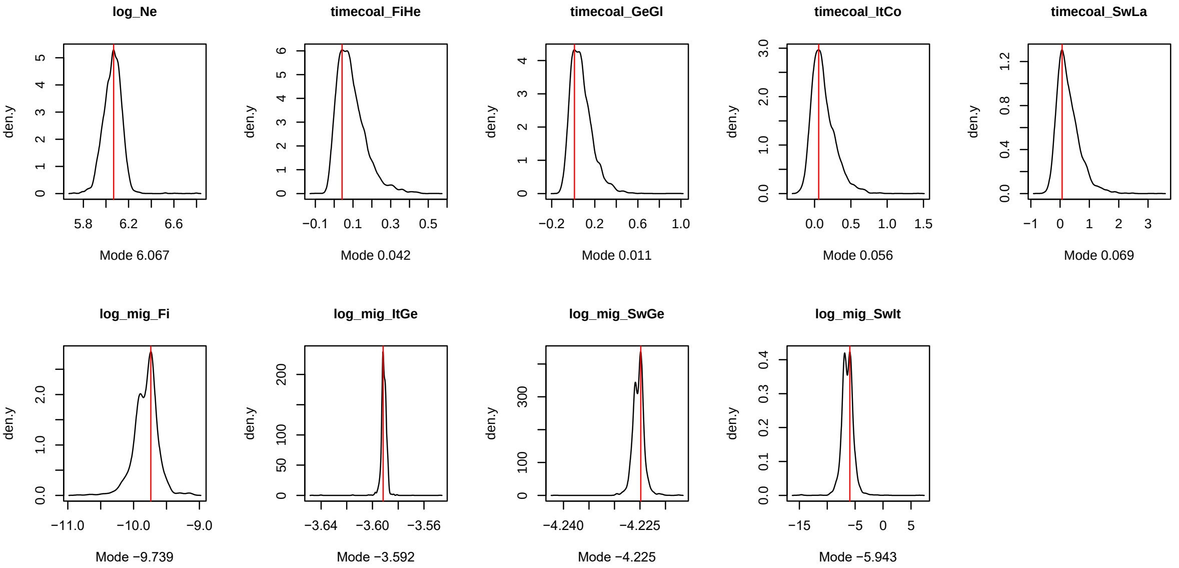


**Figure S9.** Posterior distribution of parameter estimates of Model 1. An explanation of the parameters and the characteristics of each distribution is detailed in Table 4.

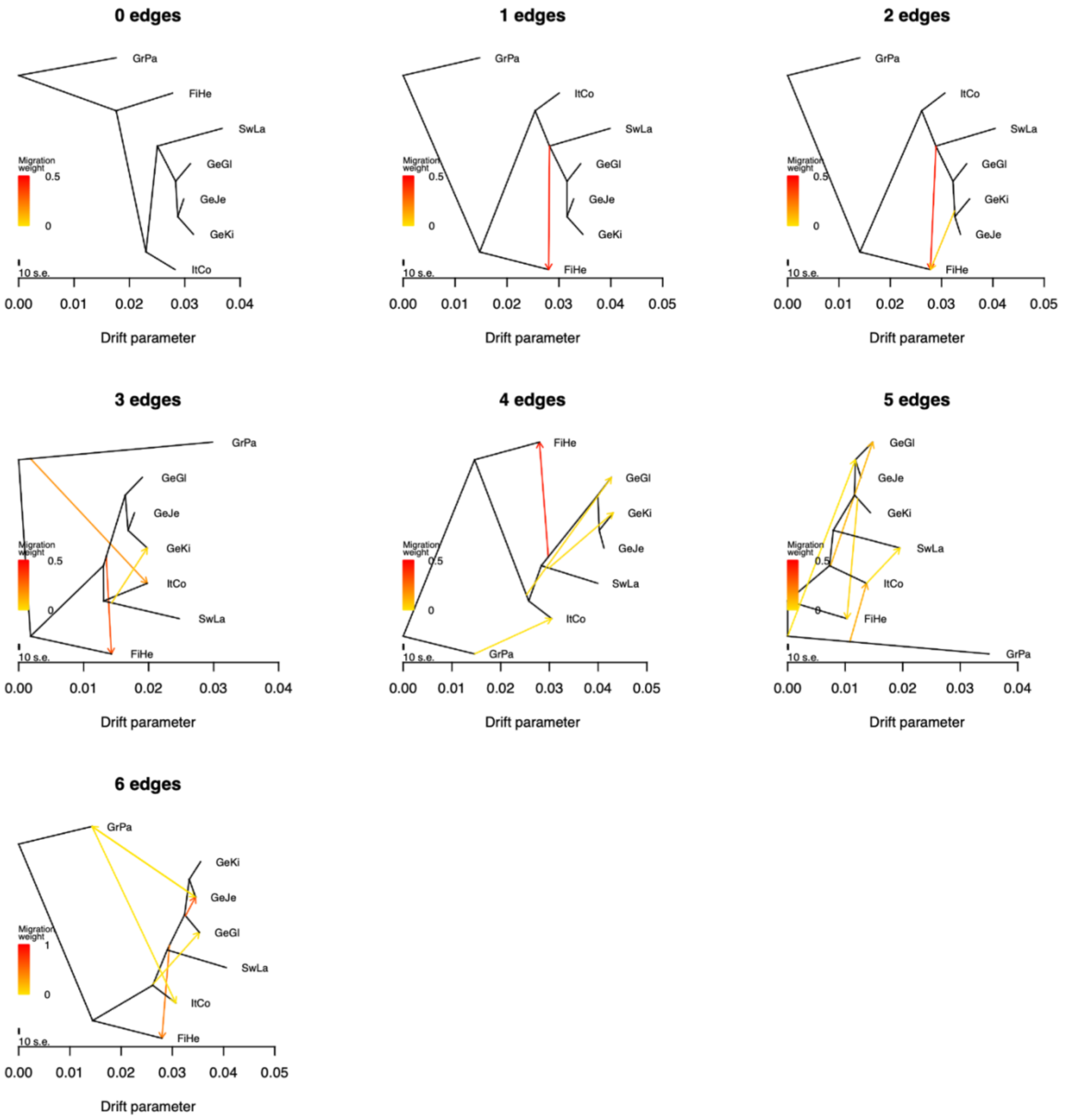


**Figure S10.** Analysis of migration between *Lampyris noctiluca* populations using Treemix. Up to six migration events were explored. Arrows show the direction of migration within the phylogenetic tree. The probability of the migration weight is shown as a gradient bar.
